# Induction of antiviral gene expression by cyclosporine A, but not inhibition of cyclophilin A or B, contributes to its restriction of human coronavirus 229E infection in a lung epithelial cell line

**DOI:** 10.1101/2023.07.20.549884

**Authors:** John E. Mamatis, Carla E. Gallardo-Flores, Taylor Walsh, Ujjwal Sangwan, Che C. Colpitts

## Abstract

The development of antivirals with an extended spectrum of activity is an attractive possibility to protect against future emerging coronaviruses (CoVs). Cyclosporine A (CsA), a clinically approved immunosuppressive drug, has established antiviral activity against diverse unrelated viruses, including several CoVs. However, its antiviral mechanisms of action against CoV infection have remained elusive, precluding the rational design of non-immunosuppressive derivatives with improved antiviral activities. In this study, we evaluated the mechanisms of CsA against HCoV-229E infection in a human lung epithelial cell line. We demonstrate that the antiviral activity of CsA against HCoV-229E is independent of classical CsA target proteins, cyclophilin A or B, which are not required host factors for HCoV-229E in A549 cells. Instead, CsA treatment induces expression of antiviral genes in a manner dependent on interferon regulatory factor 1, but independent of classical interferon responses, which contributes to its inhibitory effect against HCoV-229E infection. Our results also point to a role for the HCoV-229E nucleoprotein in antagonizing activation of type I interferon, but we show that CsA treatment does not affect evasion of innate immune signaling pathways by HCoV-229E. Overall, our findings further the understanding of the antiviral mechanisms of CsA against CoV infection and highlight a novel immunomodulatory strategy to inhibit CoV infection that may inform future drug development efforts.

## 1. Introduction

The emergence of severe acute respiratory syndrome coronavirus 2 (SARS-CoV-2) in 2019 highlights the threat posed by emerging coronaviruses (CoVs). CoVs are zoonotic, enveloped, positive-sense single-stranded RNA viruses^1, 2^ with the potential for cross-species transmission that could result in future emergence events, making the development of extended-spectrum antivirals a priority. Cyclosporine A (CsA), a widely used cyclophilin inhibitor, has broad antiviral activity and inhibits the replication of several CoVs.^3–9^ While CsA has immunosuppressive properties through suppression of T cell proliferation,^10^ a retrospective clinical study in patients with severe SARS-CoV-2 infection nonetheless showed that mortality was significantly decreased amongst patients administered CsA.^11^ Importantly, several non-immunosuppressive derivatives of CsA are available and demonstrate antiviral activity against CoV infection.^9, 12, 13^ However, the antiviral mechanisms of CsA and its derivatives against CoV infection have remained elusive. To address this gap, and to improve the potential for developing effective CsA-like molecules as CoV antivirals, we characterized the antiviral mechanisms of CsA against CoV infection, using HCoV-229E as a model.

CsA classically targets cyclophilins (Cyps), a group of peptidyl-prolyl cis/trans isomerases (PPIases) with diverse functions.^14^ While Cyps are cofactors for several viruses, their potential roles in CoV infection have remained unclear. CypA, an abundant cytoplasmic Cyp that is the classical target of CsA, interacts with the nucleocapsid (N) proteins of both SARS-CoV^15^ and human coronavirus 229E (HCoV-229E).^5^ CoV N proteins antagonize interferon (IFN) activation and signalling^16–18^ to dampen expression of interferon-stimulated genes (ISGs). CsA treatment disrupts the interaction of CypA with HCoV-229E N,^5^ highlighting a potential role for CsA in circumventing viral innate immune evasion to enhance antiviral gene expression. CypA has been implicated in innate immune evasion by multiple viruses,^19^ including hepatitis C virus (HCV),^20–22^ human immunodeficiency virus (HIV),^23, 24^ and equine arteritis virus,^25^ by enhancing replication organelle formation or capsid stability. However, potential roles for Cyps in CoV immune evasion have not yet been evaluated. We hypothesized that CoVs similarly manipulate host Cyps to antagonize cell-intrinsic antiviral pathways.

Here, we show that the antiviral activity of CsA against HCoV-229E in the A549 lung epithelial cell line does not result from its inhibition of CypA or CypB. We demonstrate that HCoV-229E, like other CoVs, suppresses activation of innate immune responses by targeting interferon regulatory factor 3 (IRF3), but that this immune antagonism is not affected by CsA treatment and thus not dependent on host Cyps. We show that CsA treatment directly induces expression of a subset of antiviral genes, which is not mediated by classical type I interferon (IFN) signalling but is dependent on the transcription factor IRF1. Overall, these findings provide new insight into antiviral immune evasion by HCoV-229E and further our understanding of the anti-CoV mechanisms of CsA, which may aid the development of novel CsA-like drugs to prepare for future emerging CoVs.

## 2. Materials and Methods

### 2.1 Cell lines and viruses

Huh7 cells (JCRB0403) were obtained from the Japanese Collection of Research Bioresources Cell Bank. 293T/17 (CRL-11268) cells were acquired from American Type Culture Collection (ATCC). Huh7 and 293T/17 cells were cultured in Dulbecco’s Modified Eagle Medium (DMEM) containing 10% fetal bovine serum (FBS) and 100 U/mL penicillin/streptomycin (complete DMEM). A549 cells (NR-52268) were obtained from BEI Resources and were cultured in Hams F-12K (Kaighn’s) medium supplemented with 10% FBS and 100 U/mL penicillin/streptomycin. A549-Dual reporter cells were obtained from Invivogen and were cultured according to manufacturer instructions. All cells were cultured at 37°C in the presence of 5% CO_2_. HCoV-229E (NR-52726) was obtained from BEI Resources and propagated on Huh7 cells as described.^26^

### 2.2 Plasmids and cloning

The IFN-β-FLuc promoter reporter plasmid and thymidine kinase (TK)-RLuc reporter plasmid were provided by Dr. Greg Towers (University College London, UK). Plasmids encoding HCoV-229E and SARS-CoV-2 N proteins were obtained from Addgene (#151912 and #141391). C-terminally FLAG-tagged 229E N and SARS-CoV-2 N constructs were cloned into pcDNA3.1 using specific primer sequences **(Supplementary Table S1)**.

For generation of stable cell lines, psPAX2 (#12260), VSV-G (# 8454) and pLentiCRISPRv2 (#52961) were obtained from Addgene. Oligos were synthesized encoding guide RNAs targeting IRF1 **(Supplementary Table S1)** and were cloned into pLentiCRISPRv2 as described.^27^ For CRISPR/Cas9-mediated knockout (KO) of CypA and CypB, or stable silencing of CypA and CypB expression by shRNA, we used plasmid constructs that we previously generated.^22^

### 2.3 Inhibitors

CsA (J63191) was purchased from Alfa Aesar. Ruxolitinib (TLRL-RUX) was obtained from Invivogen.

### 2.4 Antibodies

Antibodies against CypA (Abcam ab126738, dilution 1:5000), CypB (Abcam ab16045, dilution 1:1000), IRF1 (Thermo Scientific MA5-31996, dilution 1:1000), α-tubulin (Thermo Scientific MA5-31466, dilution 1:2500) and GAPDH (Thermo Scientific MA5-15738, dilution 1:1000) were used for western blotting. The secondary goat anti-rabbit IgG polyclonal antibody (IRDye® 800CW, LI-COR Biosciences) and goat anti-mouse IgG polyclonal antibody (IRDye® 680RD, LI-COR Biosciences) were also used.

### 2.5 Plaque assay

Huh7 cells were plated in 12-well plates at a density of 3.5 x 10^5^ cells/well in complete DMEM. HCoV-229E-containing supernatants were serially diluted in serum-free DMEM. Cells were then inoculated with serially diluted virus at 33°C for 2 hours. Inocula were removed and replaced with plaquing media (DMEM containing 2% FBS and 1.2% carboxymethylcellulose). Following 96 h at 33°C, cells were fixed and stained by addition of 0.5% crystal violet in 20% methanol to each well to visualize plaques.

### 2.6 RT-qPCR

Cellular RNA was extracted from cell lysates using the Monarch Total RNA Miniprep Kit (New England Biolabs) according to the manufacturer’s protocol. Isolated RNA was reverse transcribed to generate cDNA using the High-Capacity cDNA Reverse Transcription Kit (Thermo Scientific), and qPCR was subsequently performed using the PowerTrack™ SYBR Green Master Mix (Thermo Scientific), according to the manufacturer instructions, with specific primers **(Supplementary Table S1)**. Gene expression was normalized to cellular actin using the 2^-dCt method.

### 2.7 Generation of stable cell lines

293T/17 cells were seeded in 6-well plates at a density of 8 x 10^5^ cells/well and incubated overnight. Transfection mixes were prepared containing 800 ng of LentiCRISPRv2 or pLKO.1 transfer plasmid, 600 ng of psPAX2 packaging plasmid, 600 ng VSV-G envelope plasmid and 6 μL of Lipofectamine 2000 (Thermo Scientific). Supernatants were collected at 48 and 72 h post-transfection and clarified using 0.45 μm filters. Lentiviruses were aliquoted and stored at -80°C.

To generate shCypA and shCypB knockdown cell lines and IRF1 KO cell lines, A549 cells were seeded in 6-well plates at a density of 2.5 x 10^5^ cells/well and transduced with lentivirus in the presence of 8 μg/mL polybrene. Cells were selected in the presence of 1 μg/mL puromycin starting at 72 h post-transduction. For generation of CypA and CypB KO cell lines, pLentiCRISPRv2 was electroporated into A549 cells using the Neon Transfection System (Invitrogen) (30 ms at 1200 V for 2 pulses) as described previously.^22^ Clonal CRISPR KO cell lines were generated by limiting dilution in 96-well plates.

### 2.8 Western blot

Samples were separated by SDS-PAGE as described previously^22^ and transferred to nitrocellulose membranes using the TransBlot Turbo System (Bio-Rad) according to manufacturer instructions (25 V, 1.0 A for 30 minutes). Membranes were blocked in 1X TBS-T (Tris buffered saline with 0.1% Tween-20) containing 5% milk for 1 hour. Primary antibodies were diluted in blocking buffer and membranes were incubated at 4°C overnight. Membranes were washed and then incubated for 1 h at room temperature with Licor IRDye-680 (red) goat anti-mouse and IRDye-800 (green) goat anti-rabbit secondary antibodies diluted 1:10,000. Membranes were washed and visualized using a Licor Odyssey^®^ Clx imager.

### 2.9 Luciferase reporter assays

293T/17 cells were seeded in 24-well plates at a density of 2 x 10^5^ cells/mL in complete DMEM. The following day, cells were transfected with 100 ng of either pcDNA-SARS2-N-FLAG, pcDNA-229E-N-FLAG, or pcDNA control, together with 10 ng of IFNβ-FLuc and 5 ng of TK-RLuc, using Lipofectamine 2000. After 24 h incubation at 37°C, cells were stimulated by transfection with 1 μg of poly(I:C) (tlrl-pic; Invivogen) for an additional 8 hours at 37°C. Cells were then lysed with 100 μL/well of 1X passive lysis buffer (Promega) and lysates (20 μL/well) were transferred into two separate 96-well white plates. For firefly luciferase readings, 20 μL of lysate was incubated at room temperature with 20 μL of BrightGlo reagent (Promega) for 5 minutes. For measurement of Renilla luciferase activity, 50 μL of prepared 1 mg/mL coelenterazine substrate (NanoLight) was injected into each well. Luciferase activity was measured using a GloMax plate reader, and data were normalized by determining the ratio of FLuc:RLuc signal.

### 2.10 A549-Dual reporter assays

A549-Dual reporter cells were seeded in 12-well plates at a density of 2.5 x 10^5^ cells/mL and incubated overnight at 37°C. Cells were mock-infected or infected with HCoV-229E, and treated with CsA or DMSO. As controls, cells were transfected with 4 μg of poly(I:C) to activate IRF3, or treated with 100 ng/mL TNF-α (rcyc-htnfa; Invivogen) to activate NF-κB. QUANTI-Luc and QUANTI-Blue assays (Invivogen) to assess IRF3 or NF-κB activation, respectively, were performed according to the manufacturer’s instructions.

### 2.11 siRNA transfection

To silence IRF1 expression, 10 μM of siRNA targeting IRF1 (siIRF1; Thermo #115266) or a control siRNA (siCTRL; Thermo #AM4611) were diluted with 3 μL of Lipofectamine RNAiMAX in Opti-MEM and incubated at room temperature for 15 minutes. Reverse transfection in A549 cells was performed according to the manufacturer’s instructions.

### 2.12 Cell viability assay

A549 cells were seeded at 1 or 2 x 10^4^ cells/well in a 96-well plate in complete F12K. After 24 h, cells were treated with serially diluted CsA for 48 h at 37°C. Cell viability was assessed using the alamarBlue^TM^ reagent (Thermo Scientific), according to the manufacturer’s instructions, using a SpectraMax® iD3 Multi-Mode Microplate Reader.

### 2.13 Statistical Analysis

Statistical analyses were conducted using GraphPad Prism 9 version 9.1.3.

## 3. Results

### 3.1 HCoV-229E infection is inhibited by CsA treatment, but not CypA or CypB depletion, in A549 lung epithelial cells

Previous studies have demonstrated that CsA and non-immunosuppressive CsA derivatives inhibit HCoV-229E infection, by unclear mechanisms, in Huh7 and Huh7-derived human hepatoma cell lines.^3–5^ Here, we evaluated the antiviral mechanisms of CsA against HCoV-229E infection in A549 human lung alveolar epithelial cells. We first infected A549 cells with HCoV-229E at a multiplicity of infection (MOI) of 0.01 plaque-forming units (pfu)/cell. After 2 h infection, inocula were removed and cells were treated with 10 μM CsA or dimethyl sulfoxide (DMSO) vehicle control for 48 h. Consistent with previous findings in Huh7-derived cells,^3–5^ CsA treatment decreased HCoV-229E titer **(Figure 1A)**. To determine the half-maximal inhibitory concentration (IC_50_), cells were infected with HCoV-229E and treated with increasing concentrations of CsA ranging from 0.05 to 10 μM. Supernatants were collected to determine viral titer, and data was expressed as a percentage relative to DMSO control. CsA dose-dependently inhibited HCoV-229E infection (IC_50,_ 0.15 μM) without affecting cell viability under these conditions **(Figure 1B)**, although we noted a minor effect on cell viability at the highest CsA concentrations when cells were seeded at a sub-confluent density **(Supplementary Fig. S1)**. Consistent with the effect on viral titer, HCoV-229E N gene expression **(Figure 1C)** was decreased by 100-fold in CsA-treated cells relative to DMSO-treated cells, with a corresponding profound reduction in N protein expression as assessed by western blot **(Figure 1D)**.

**Figure 1.**
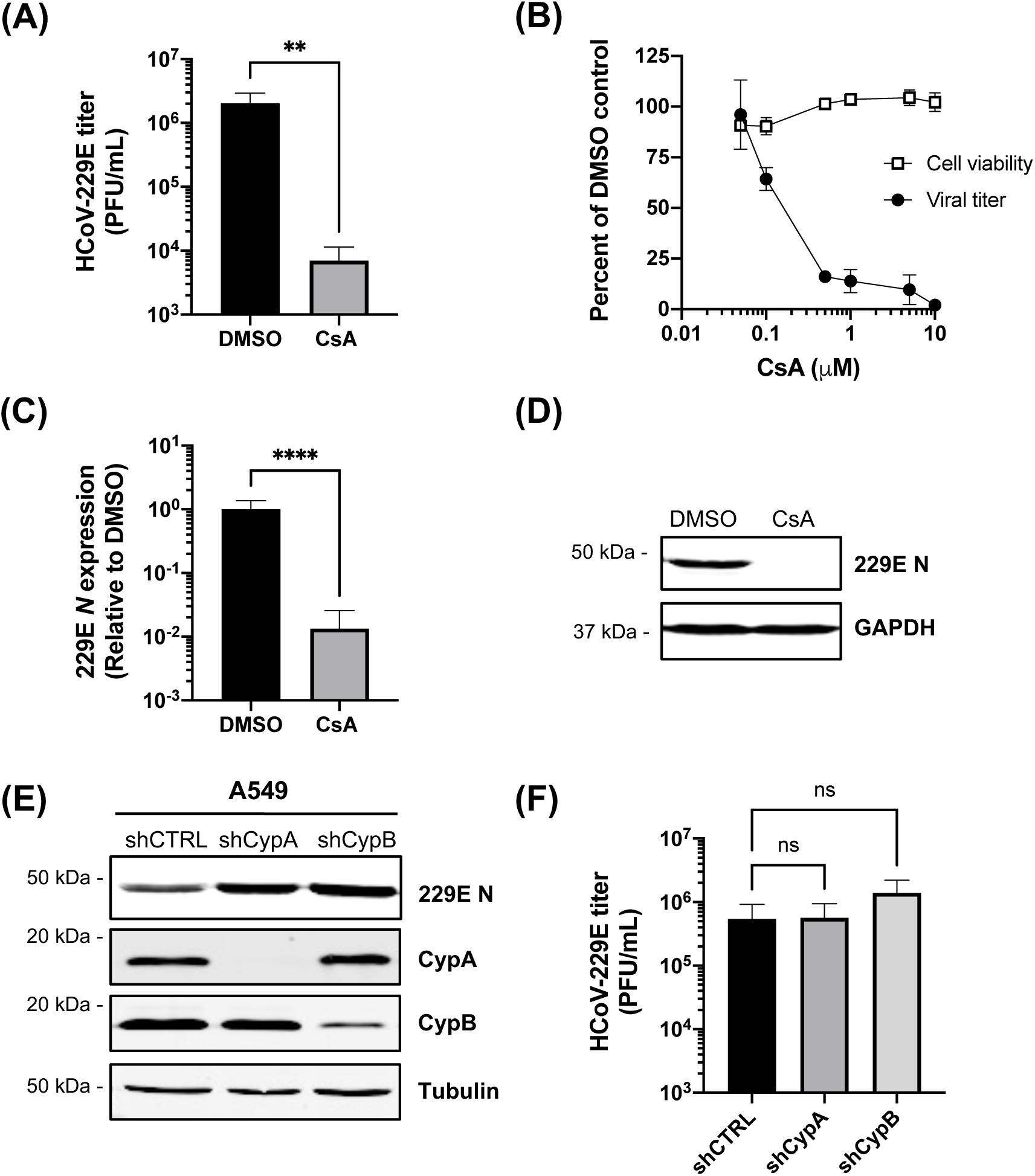
Replication of HCoV-229E in A549 cells is inhibited by CsA treatment, but not CypA or CypB depletion. **(A-D)** CsA treatment inhibits HCoV-229E replication in A549 cells. A549 cells were infected with HCoV-229E (MOI 0.01) and treated with 10 μM CsA or DMSO vehicle control for 48 h. **(A)** The viral titer following 10 μM CsA treatment was measured by plaque assay. **(B)** Dose-response analysis of the effect of CsA treatment on HCoV-229E viral titer. The viral titer was quantified by plaque assay and cell viability was assessed using the alamarBlue reagent. **(C-D)** HCoV-229E *N* gene expression was measured by RT-qPCR (normalized to actin) and HCoV-229E N protein expression was determined by western blot. **(E-F)** HCoV-229E replication is not affected by CypA or CypB depletion in A549 cells. Western blot assessing HCoV-229E N protein, CypA or CypB expression in A549 cells transduced with shRNAs targeting CypA or CypB. **(F)** Plaque assay quantifying HCoV-229E viral titer following infection of CypA and CypB knockdown cell lines. Cells were infected at MOI 0.01 and incubated for 48 h. Graphs show the average +/- standard deviation (SD) from 3 independent experiments, with qPCR **(C)** performed in technical triplicates. Significance was determined by unpaired t-test with Welch’s correction **(A, C)** or ordinary one-way ANOVA **(F)** (****p<0.0001; **p<0.01; ns, not significant).

To evaluate potential Cyp host factor dependencies, we stably silenced expression of CypA or CypB in A549 cells using shRNA **(Figure 1E)** and evaluated HCoV-229E replication. Neither HCoV-229E nucleoprotein expression nor viral titer were significantly affected by CypA or CypB depletion **(Figure 1E-F)**, leading us to conclude that that the antiviral effect of CsA against HCoV-229E infection in A549 cells **(Figure 1A-D)** is not the result of CypA or CypB inhibition. Interestingly, silencing of CypA expression in Huh7 cells **(Supplementary Fig. S2A)** significantly decreased HCoV-229E *N* gene expression **(Supplementary Fig. S2B)** and viral titer **(Supplementary Fig. S2C)**, reflecting a previous study in Huh7-derived cells^28^ and highlighting cell type-dependent Cyp requirements. However, we focused our subsequent efforts on characterizing the antiviral mechanism of CsA against HCoV-229E infection in A549 cells, as these cells are more physiologically relevant for CoV infection.

### 3.2 CsA induces antiviral gene expression in HCoV-229E-infected cells independently of CypA or CypB

CsA treatment induces ISG expression during HCV,^22, 29, 30^ rotavirus,^31, 32^ HIV,^23^ and MERS-CoV infection.^8^ Therefore, we evaluated whether CsA similarly enhances antiviral gene expression in HCoV-229E-infected A549 cells, which may contribute to its antiviral effect. HCoV-229E-infected cells were treated with DMSO or CsA for 48 h, at which point cell lysates were collected for RT-qPCR analysis. Interestingly, CsA treatment induced expression of a representative ISG, *MX1*, in HCoV-229E-infected A549 and Huh7 cells **(Figure 2A)**. We next performed a time-course experiment to evaluate the kinetics of *MX1* induction following CsA treatment of A549 cells infected with HCoV-229E (MOI 0.05). *MX1* expression was significantly increased in CsA-treated cells relative to the DMSO-treated control cells at 48 and 72 h post-infection (hpi) **(Figure 2B)**. A similar phenotype was observed when cells were infected at a higher MOI (0.5) and treated similarly with CsA **(Supplementary Fig. S3)**.

**Figure 2.**
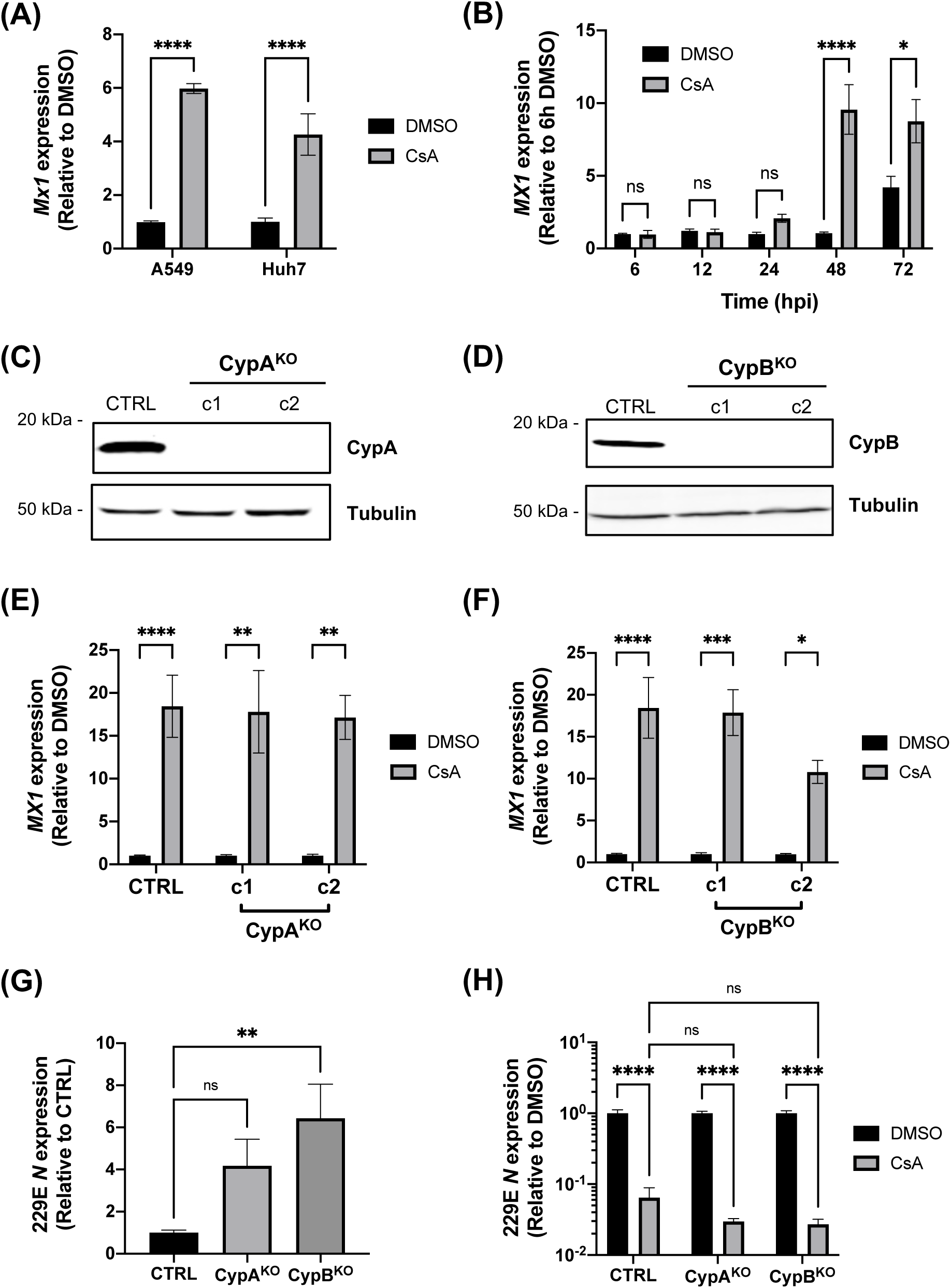
Induction of antiviral gene expression by CsA is not mediated by CypA or CypB. **(A)** CsA treatment induces ISG expression in A549 and Huh7 cells. A549 cells or Huh7 cells were infected with HCoV-229E (MOI 0.01) and treated with 10 μM CsA or DMSO vehicle. Cell lysates were collected after 48 hours and *MX1* expression was determined by RT-qPCR (normalized to actin). **(B)** CsA induces *MX1* expression at 48 and 72 hpi, but not earlier time points, in HCoV-229E-infected cells. A549 cells were infected with HCoV-229E (MOI 0.01) and treated or not with CsA. Cell lysates were collected at indicated time points to evaluate *MX1* expression by RT-qPCR. **(C-D)** Western blot demonstrating successful knockout of CypA **(C)** and CypB **(D)** in A549 cells. A549 CypA KO cells **(E)** or A549 CypB KO cells **(F)** were infected with HCoV-229E (MOI 0.01) and treated with 10 μM CsA or DMSO. Expression of *MX1* at 48 hpi was evaluated by RT-qPCR. Graphs show the average +/- standard error of the mean (SEM) from 3 independent experiments, each performed in technical triplicates. Statistical significance was assessed by two-way ANOVA (****p<0.0001; ***p <0.001; **p<0.01; *p<0.05; ns, not significant). **(G-H)** A549 CypA^KO^ cells (c1) or CypB^KO^ cells (c1) were infected with HCoV-229E (MOI 0.01) and left untreated **(G)** or treated with 10 μM CsA or DMSO **(H)**. At 48 hpi, HCoV-229E N gene expression was assessed by RT-qPCR. Graphs show the average +/- standard error of the mean (SEM) from 2 independent experiments, each performed in technical triplicates. Statistical significance was assessed by one-way ANOVA (A) or two-way ANOVA (B) ****p<0.0001; **p<0.01; ns, not significant).

CypA regulates the ubiquitination of RIG-I^33^ and has also been implicated in NF-κB activation,^34^ while CypB was shown to regulate IRF3 activation.^35^ Since NF-κB and IRF3 are required for the production of type I IFN and subsequent ISG expression during RNA virus infection, we evaluated potential roles for Cyps in modulating the ISG induction observed in CsA-treated cells. To avoid potential immunostimulatory effects of shRNA, we generated clonal A549 CypA or CypB KO cell lines using CRISPR/Cas9 **(Figure 2C-D)**. CsA still induced *MX1* expression in the absence of CypA or CypB **(Figure 2E-F)**, demonstrating that neither Cyp is required for CsA-induced ISG expression. Furthermore, viral *N* gene expression was not reduced in the CypA or CypB KO cell lines **(Figure 2G)**, and CsA treatment inhibited HCoV-229E replication in CypA or CypB KO cells **(Figure 2H)**, supporting our conclusion that CsA inhibits HCoV-229E replication independently of CypA or CypB in A549 cells.

### 3.3 CsA does not impact HCoV-229E evasion of IFN

The interaction between HCoV-229E N protein and CypA identified in Huh7 cells was disrupted by CsA treatment,^5^ which targets the conserved PPIase active site of CypA, suggesting that other Cyps could similarly interact with 229E N protein via their PPIase domains. Since previous literature has demonstrated that the SARS-CoV-2 nucleoprotein (N) represses IFN-β reporter activity,^36^ we hypothesized that CsA may interfere with Cyp-N interactions to disrupt N-mediated immune evasion, restoring ISG expression in infected cells and explaining the increased ISG expression we observed **(Figure 2A-B)**. To test this hypothesis, we evaluated the effect of CsA treatment on N-mediated antagonism of IFN-β activation by luciferase reporter assay. We cloned HCoV-229E N or SARS-CoV-2 N into pcDNA3.1 to generate C-terminally FLAG-tagged pcDNA-229E-N or pcDNA-SARS2-N. These plasmids were transfected in HEK293T/17 cells **(Figure 3A)** along with an IFN-β-responsive promoter reporter plasmid driving firefly luciferase expression and a thymidine kinase-responsive promoter reporter plasmid driving Renilla luciferase expression. Activation of IFN-β was assessed by measuring firefly luciferase activity normalized to Renilla luciferase activity from cell lysates. Consistent with published literature,^36^ transfection of SARS-CoV-2 N inhibited IFN-β reporter activity, and we identified a similar inhibitory effect for HCoV-229E N **(Figure 3B)**. However, CsA treatment did not affect the inhibition of IFN-β reporter activity by SARS-CoV-2 or HCoV-229E N protein **(Figure 3B)**.

**Figure 3.**
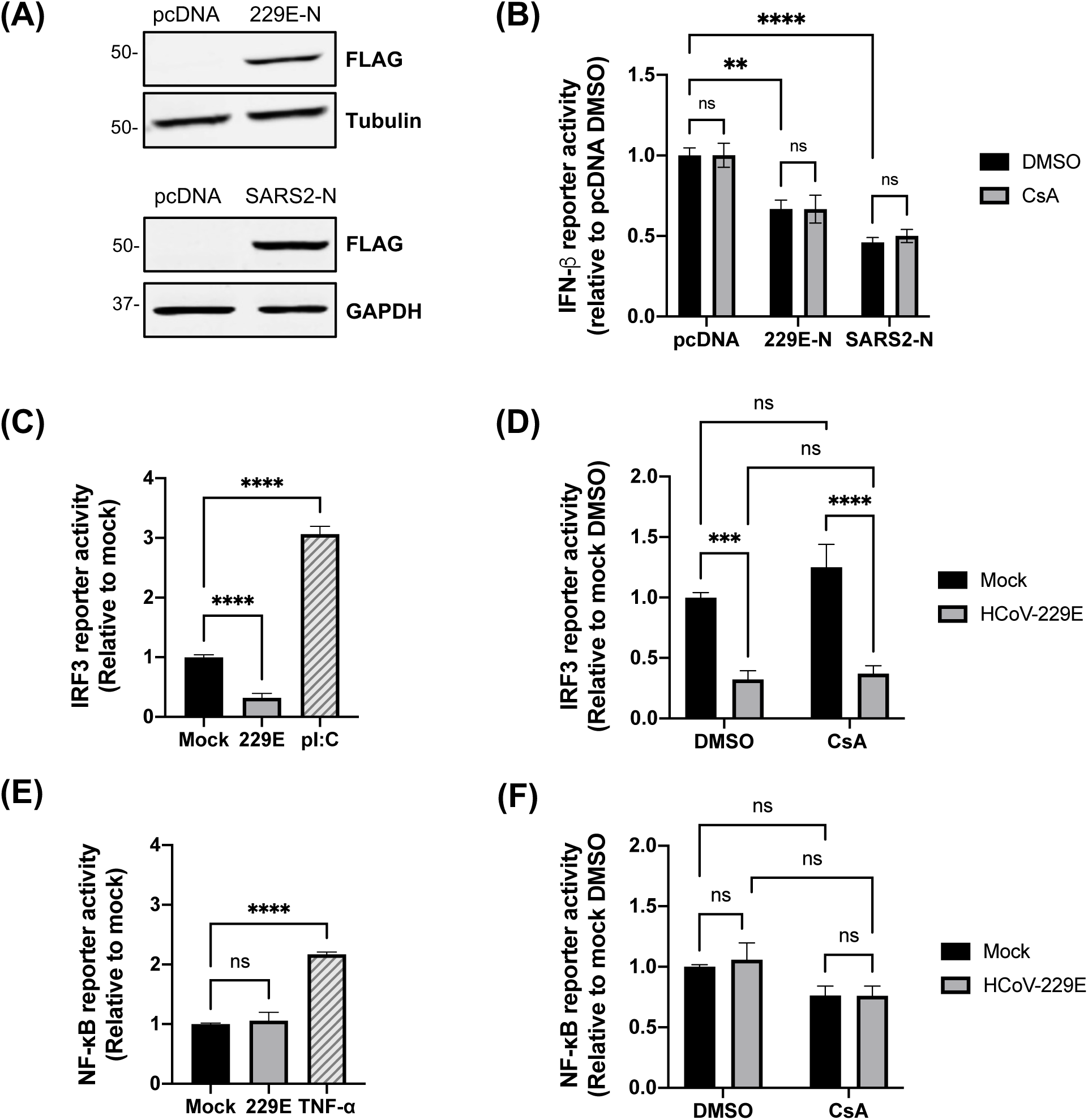
CsA treatment does not impact HCoV-229E evasion of IFN responses. **(A)** HEK-293T/17 cells were transfected with pcDNA empty vector control or expression plasmids encoding FLAG tagged HCoV-229E N (229E-N) or SARS-CoV-2 N (SARS2-N) for 24 hours. FLAG expression was confirmed by western blot at the expected molecular weight of the respective N proteins. **(B)** Ectopic expression of CoV N protein inhibits IFN activation. HEK293T/17 cells were co-transfected with 100 ng of either 229E N, SARS2 N, or pcDNA empty vector control, together with 10 ng of IFN-β firefly luciferase reporter and 5 ng of thymidine kinase Renilla luciferase reporter control constructs. After 24 hours in the presence of 10 μM CsA or DMSO, cells were stimulated with poly(I:C) for 8 hours and subsequently lysed. Luciferase reporter activity was measured from cell lysates. Data are normalized as a ratio of FLuc:RLuc, relative to pcDNA empty vector control. **(C-F)** A549 Dual reporter cells were mock-infected or infected with HCoV-229E (MOI 0.01) for 48 hours. **(C-D)** IRF3 activation was assessed by quantification of Lucia luciferase reporter activity, using poly(I:C) (4 μg) as a positive control to activate IRF3 **(C)**. HCoV-229E suppressed IRF3 reporter activity **(C)** in the presence or absence of 10 μM CsA **(D)**. **(E-F)** NF-κB activation was assessed by quantification of SEAP activity in cell supernatants, using TNF-α (100 ng/mL) as a positive control to active NF-κB **(E)**. Mock infected and HCoV-229E-infected A549 Dual cells were treated with 10 μM CsA or DMSO vehicle control for 48 hours. Neither HCoV-229E infection (E) nor CsA treatment (F) significantly affected NF-κB reporter activity. Graphs show the average +/-SEM from 3 independent experiments. Statistical significance was tested by one-way ANOVA **(C, D)** or two-way ANOVA **(B, D, F)** (****p<0.0001; ***p<0.001; **p<0.01; ns, not significant).

CoVs have evolved a multitude of mechanisms to inhibit IFN activation.^37^ To more broadly evaluate whether CsA disrupts antagonism of type I IFN by HCoV-229E, we used A549-Dual reporter cells, which express inducible reporters to measure IRF3 or NF-κB activity, to probe IFN activation during HCoV-229E infection. IRF3 reporter activity was inhibited in HCoV-229E-infected cells relative to mock-infected cells **(Figure 3C)**, consistent with antagonism of IRF3 by HCoV-229E. However, reflecting our IFN-β reporter assay results **(Figure 3B)**, CsA treatment did not affect modulation of IRF3 reporter activity in HCoV-229E-infected cells **(Figure 3D)**. Since coordinated activation of IRF3 and NF-κB is required for production of IFN-β,^38, 39^ we also tested the effects of HCoV-229E infection and CsA treatment on NF-κB activation. NF-κB reporter activity was not significantly affected by HCoV-229E infection **(Figure 3E)** or CsA treatment **(Figure 3F).** Taken together, these findings show that HCoV-229E primarily evades type I IFN activation by targeting IRF3, and this occurs independently of host Cyps.

### 3.4 CsA induces expression of a subset of ISGs independently of classical IFN signalling

Although CsA treatment did not significantly affect IRF3 or NF-κB activation in mock-infected cells **(Figure 3D, 3F)**, we noted that CsA treatment induced *MX1* expression in uninfected A549 cells after 48 hours **(Figure 4A)**, albeit to a lesser extent than observed in infected cells. Thus, CsA directly induces ISG expression, but this occurs independently of the classical IFN activation pathway mediated by IRF3 and NF-κB. To further probe the role of IFN, we evaluated whether CsA-induced ISG expression requires IFN signalling, using ruxolitinib to antagonize IFN signalling. Ruxolitinib suppresses type I and type III IFN signalling by preventing activation of JAK/STAT signalling and subsequent ISG induction.^40^ As expected, ruxolitinib blocked the ability of IFN-β to inhibit HCoV-229E infection **(Supplementary Fig. S4)**. Cells were infected with HCoV-229E and treated with CsA or DMSO vehicle in the presence or absence of ruxolitinib. The induction of *MX1* expression by CsA was not affected by ruxolitinib treatment **(Figure 4B)**, nor did ruxolitinib significantly affect the ability of CsA to inhibit HCoV-229E infection, as measured by viral titer **(Figure 4C)** or *N* gene expression **(Figure 4D)**. To gain further insight into the mechanism underlying ISG induction by CsA, we evaluated whether the expression of other ISGs was similarly induced by CsA treatment. Interestingly, we observed that CsA treatment significantly induced expression of a subset of ISGs, including *MX1* and *OAS2*, but not *TLR3* or *ISG15* **(Figure 4E)**, suggesting that CsA does not broadly induce ISG expression but rather modulates specific pathways or transcription factors.

**Figure 4.**
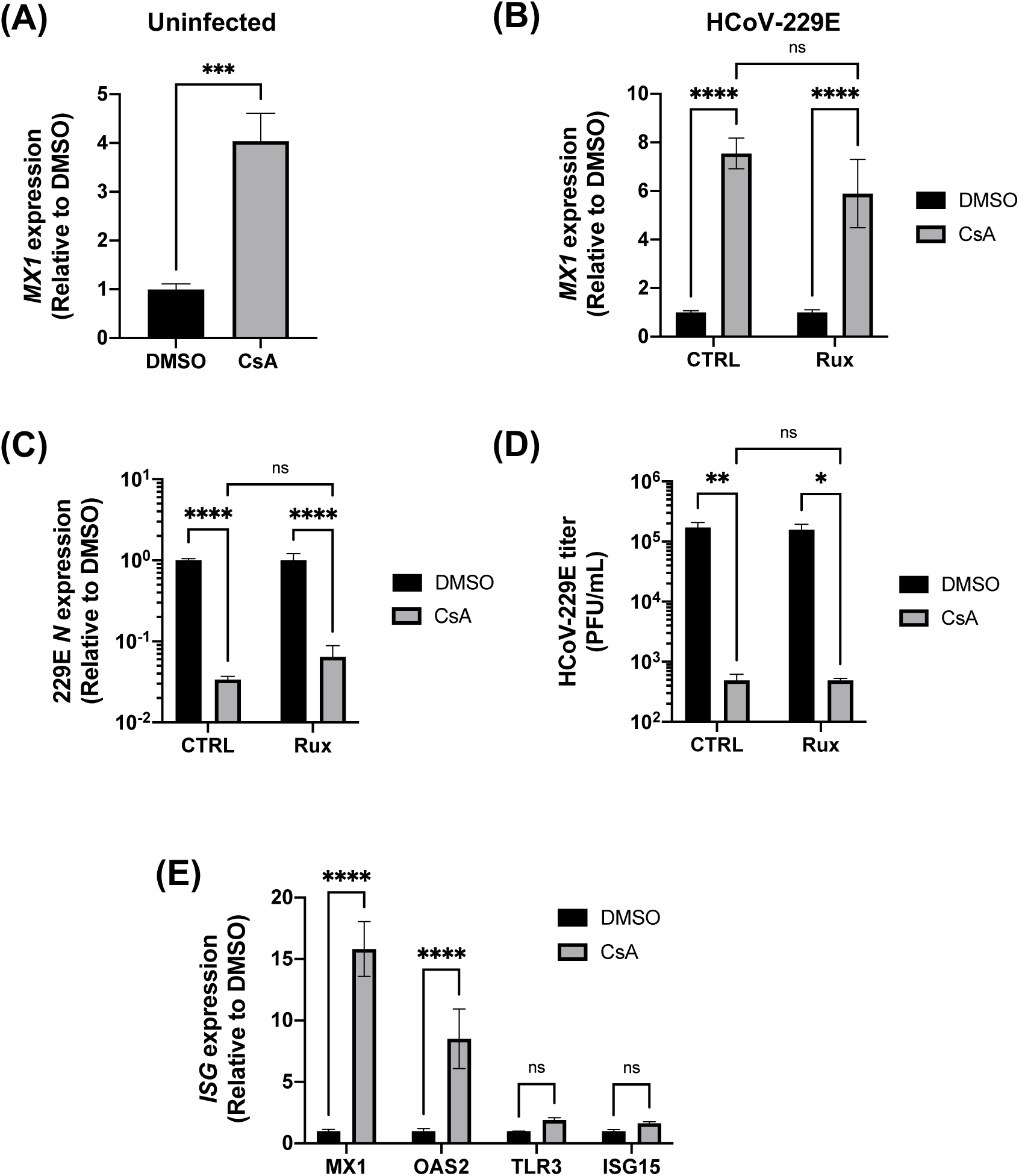
CsA induces expression of a subset of ISGs independently of classical type I IFN signaling. **(A)** Uninfected A549 cells were treated with 10 μM CsA or DMSO for 48 h, and MX1 expression was evaluated by RT-qPCR (normalized to actin). **(B-D)** A549 cells were infected with HCoV-229E (MOI 0.01) and were treated with 10 μΜ CsA or DMSO in the presence or absence of ruxolitinib (Rux) for 48 hours. Rux treatment does not affect the ability of CsA to induce *MX1* expression **(B)** or suppress viral *N* gene expression **(C)**, as determined by RT-qPCR (normalized to actin). **(D)** Viral titer was determined by plaque assay. **(E)** CsA treatment of HCoV-229E-infected A549 cells induced expression of *MX1* and *OAS2*, but not *TLR3* or *ISG15* expression, as quantified by RT-qPCR. Graphs show the average +/- SD **(D)** from 3 independent experiments, or average +/- SEM **(A-C, E)** from 3 independent experiments performed in technical triplicates. Statistical significance was assessed by unpaired t-test **(A)** or two-way ANOVA **(B-E)** (****p<0.0001; ***p <0.001; **p<0.01; *p<0.05; ns, not significant).

### 3.5 IRF1 expression modulates CsA antiviral activity and induction of ISG expression

Having ruled out a role for type I IFN in CsA-induced ISG expression, we next explored the role of IRF1, a transcription factor that has been demonstrated to drive the expression of a subset of antiviral genes, including *MX1* and *OAS2*, in an IFN-independent manner.^41, 42^ CsA has been shown to modestly induce IRF1 expression^29^ as well as expression of IRF1-dependent ISGs^22^ in HCV-infected Huh7 cells. To evaluate a role for IRF1 in our model, we used CRISPR/Cas9 to generate A549 IRF1 KO cells **(Figure 5A)**. Strikingly, CsA-induced *MX1* expression was significantly reduced in uninfected IRF1 KO cells **(Figure 5B)** and was abrogated in HCoV-229E-infected IRF1 KO cells **(Figure 5C)**, highlighting a key role for IRF1 in the induction of *MX1* expression by CsA. Furthermore, the absence of IRF1 decreased the antiviral effect of CsA against HCoV-229E infection by approximately 10-fold **(Figure 5D)**. To rule out clonal effects of the IRF1 KO cells, we transiently silenced IRF1 expression using siRNA **(Supplementary Fig. S5A)** and observed a similar phenotype **(Supplementary Fig. S5B-C)** as in the KO cells.

**Figure 5.**
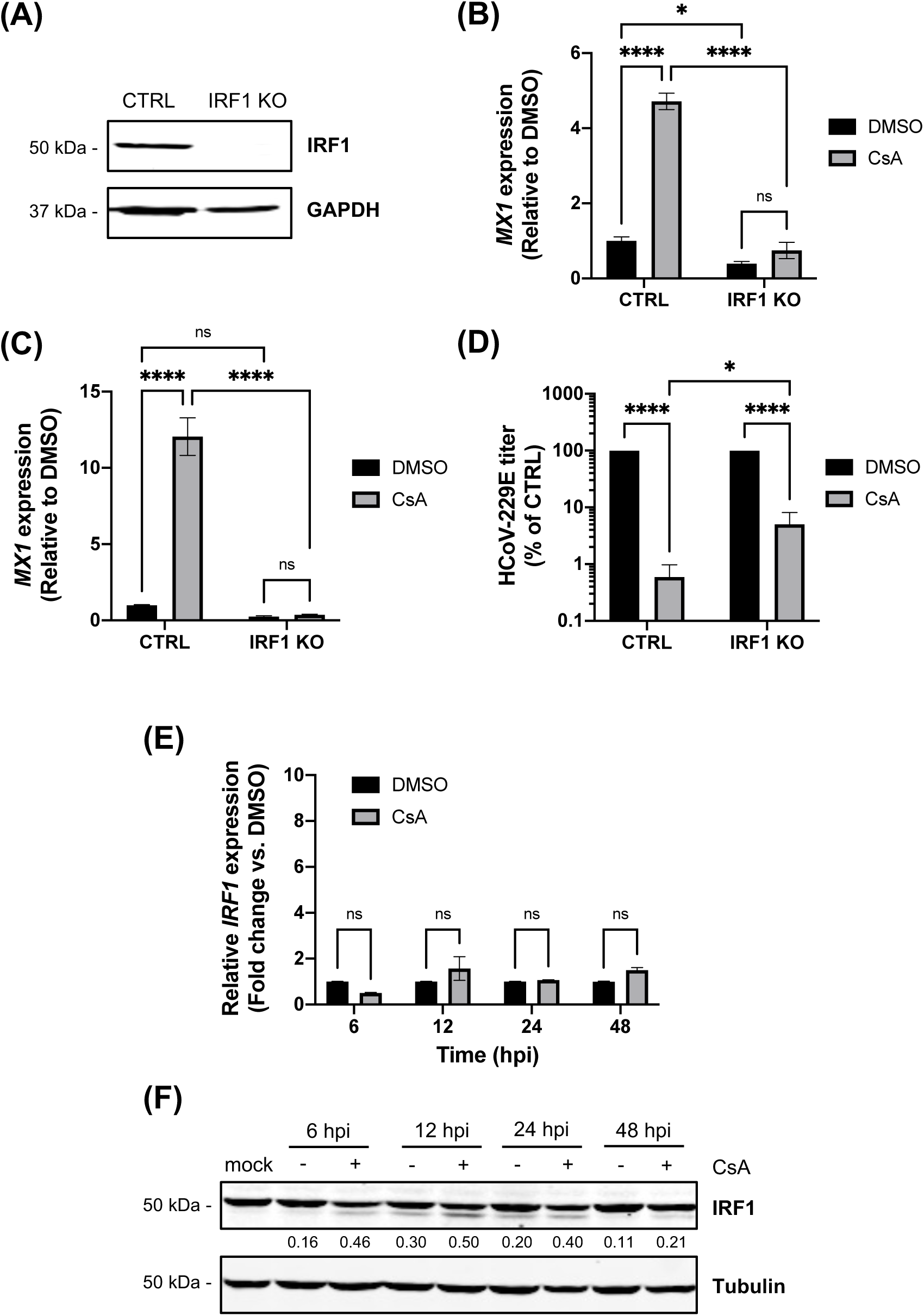
IRF1 expression is required for CsA-induced ISG expression and modulates the antiviral potency of CsA. **(A)** Western blot confirming knockout of IRF1 in A549 cells. (B-C) MX1 expression was no longer induced by CsA treatment in IRF1 KO cells. A549 IRF1 KO or control cells were either uninfected **(B)** or infected with HCoV-229E (MOI 0.01) **(C)** and treated with 10 μM CsA or DMSO control for 48 h. *MX1* expression was determined by RT-qPCR (normalized to actin). **(D)** The antiviral effect of CsA was reduced in IRF1 KO cells. Cells were infected and treated as described above, and supernatants were collected at 48 hpi to measure HCoV-229E titer by plaque assay. **(E-F)** A549 cells were infected with HCoV-229E (MOI 0.01) and treated with 10 μM CsA or DMSO vehicle for the indicated time points. IRF1 expression was evaluated by RT-qPCR **(E)** or western blot **(F)**. One representative blot is shown, with the band density ratio of the lower molecular weight IRF1 band to the higher molecular weight band indicated. Graphs show the average +/- SEM from 3 independent experiments each performed in technical triplicates **(B, C, E)**, or average +/- SD from 3 independent experiments **(D).** Statistical significance was assessed by two-way ANOVA (****p<0.0001; *p<0.05; ns, not significant).

We next evaluated whether CsA treatment increased IRF1 expression, which we hypothesized would enhance expression of IRF1 target genes such as *MX1*. Unexpectedly, IRF1 expression in A549 cells was not significantly enhanced by CsA treatment, either at the mRNA **(Figure 5E)** or protein level **(Figure 5F)**. We speculate that CsA treatment may instead affect post-translational modifications of IRF1 that modulate its activation. We consistently observed a lower molecular weight IRF1 band in samples treated with CsA that was either absent or less pronounced in DMSO-treated samples **(Figure 5F)**. We aim to perform future studies to evaluate the effects of CsA on IRF1 post-translational modifications.

## 4. Discussion

In this study, we characterized the antiviral mechanisms of CsA against the human alphacoronavirus HCoV-229E. We show that the antiviral effect of CsA against HCoV-229E is independent of CypA or CypB, but is partially dependent on IRF1, a transcription factor driving IFN-independent expression of certain ISGs.^41, 42^ Our finding that CypA and CypB are not required for HCoV-229E infection in A549 cells **(Figure 1E-F, 2G)**, but that CypA is required in Huh7 cells **(Supplementary Fig. S2)**, is consistent with published literature, where silencing of CypA expression by shRNA in Huh7.5 cells led to a decrease in HCoV-229E reporter gene expression.^28^ An interaction between CypA and HCoV-229E N protein was also identified in Huh7 cells,^5^ although de Wilde et al. showed that HCoV-229E infection in Huh7 cells was not inhibited by CRISPR/Cas9-mediated knockout of CypA, CypB, CypC or CypD,^43^ reflecting our findings in A549 cells where depletion of CypA or CypB by shRNA or CRISPR did not inhibit HCoV-229E infection **(Figure 1E-F)**. Clearly, subtle cell type or cell line differences contribute to the differential Cyp requirements observed in different studies. The underlying mechanisms remain to be explored. However, reported Cyp dependencies are inconsistent among human CoVs, where CypA is required for HCoV-NL63 infection^7^ but not for HCoV-229E or MERS-CoV infection.^43^ Similarly, CypA is not essential for SARS-CoV-2 replication in Calu-3 cells,^44^ and numerous screens for SARS-CoV-2 host factors failed to identify compelling Cyp candidates.

Since Cyp dependencies are not conserved, it is likely that an alternative mechanism, such as the induction of an antiviral state in cells, contributes to the pan-CoV and broad-spectrum antiviral effect of CsA. We previously showed that CsA treatment enhances antiviral gene expression in HCV-infected Huh7 cells.^22^ In parallel, Sauerhering et al. showed that CsA treatment induces the IRF1-IFN-α (type III IFN) signalling axis in Calu-3 cells, which contributes to its antiviral effect against MERS-CoV.^8^ Here, our data support a role for IRF1 in mediating the antiviral effect of CsA against another CoV, HCoV-229E. However, the mechanisms may be different, as blockade of type I and type III IFN signaling did not significantly impact ISG induction or the antiviral efficacy of CsA **(Figure 4B-D)**, pointing to direct induction of ISG expression by IRF1,^43^ in contrast to the central role of type III IFN observed in MERS-CoV-infected Calu-3 cells.^8^ However, further studies to more explicitly evaluate a role for type III IFN are warranted. It is also worth noting that CsA retained some antiviral activity against HCoV-229E in IRF1-depleted A549 cells, suggesting a role for additional inhibitory mechanisms yet to be determined. Nonetheless, it is likely that induction of ISG expression by CsA via the IRF1 pathway contributes to or enhances its antiviral effect against diverse viral infections.

The mechanisms underlying the CsA-induced expression of IRF1-dependent ISGs remain to be precisely determined. In Huh7^29^ and Calu-3 cells,^8^ CsA treatment modestly upregulates the expression of IRF1 (but not other IRFs) at the mRNA level. In contrast, CsA treatment does not enhance IRF1 expression in A549 cells **(Figure 5E-F)**, although IRF1 is required for CsA-induced ISG expression **(Figure 5B-C)**, suggesting that CsA may affect IRF1 activity at a post-transcriptional and post-translational level. Our study contributes new insight into the regulation of IRF1 activity by CsA. First, we show that neither CypA nor CypB are required for CsA-mediated IRF1-dependent ISG induction **(Figure 2E-F)**. Secondly, CsA induces expression of IRF1-dependent antiviral genes independently of NF-κB or IRF3 activation **(Figure 3D, 3F)**. We speculate that CsA may promote IRF1 activation by regulating its post-translational modifications, nuclear translocation, or protein complex formation. Indeed, CsA will be a useful tool for future studies exploring regulation of IRF1.

We had initially hypothesized that CsA may disrupt innate immune evasion by HCoV-229E, as has been observed for other viruses.^19^ While this did not appear to be the case, our findings nonetheless contribute new insight into HCoV-229E antagonism of IFN activation, which has not been well characterized yet. We demonstrate that HCoV-229E antagonizes IFN activation by targeting the IRF3 pathway **(Figure 3C)**, and that this is mediated, at least in part, by its N protein **(Figure 3B)**. SARS-CoV-2 N protein has been shown to antagonize RIG-I and MAVS activation.^37^ However, with only 30% sequence identity between SARS-CoV-2 N and HCoV-229E N proteins,^45^ it is possible that the HCoV-229E N protein may interfere with IRF3 activation by different mechanisms, which remain to be explored.

In conclusion, our findings contribute to understanding the antiviral mechanisms of CsA, using HCoV-229E infection in a human lung cell line as a model. Although CsA itself is not likely to be useful clinically as an antiviral drug, due to its suppression of T cell proliferation, understanding the mechanisms underlying its broad-spectrum antiviral activity will inform medicinal chemistry efforts aimed at developing non-immunosuppressive CsA derivatives suitable for further development as extended-spectrum antivirals. Such molecules may be useful to enhance preparedness for future emerging viral outbreaks.

## Declaration of Competing Interests

The authors declare that they have no known competing financial interests or personal relationships that could have appeared to influence the work reported in this paper.

## Acknowledgements

This study was supported by a discovery grant from the Natural Sciences and Engineering Research Council of Canada (RGPIN/04277-2020), a priority announcement grant from the Canadian Institutes for Health Research (184008), the Banting Research Foundation, the J.P. Bickell Foundation for Medical Research, the Canadian Foundation for Innovation John R. Evans Leaders Fund, and Queen’s University. CEGF is supported by a PhD studentship from the Canadian Network on Hepatitis C Virus. The following reagents were obtained through BEI Resources (NIAID, NIH): *Homo sapiens* A549 Lung Carcinoma Cells (NR-52268); Human Coronavirus 229E (NR-52726).

## Notes

### Competing Interest Statement

The authors have declared no competing interest.

