## Supplemental Figures for "Induction of antiviral gene expression by cyclosporine A, but not inhibition of cyclophilin A or B, contributes to its restriction of human coronavirus 229E infection in a lung epithelial cell line"

### Supplementary table

***Supplementary Table S1.*** *Oligo sequences (5' -> 3')*

| | | |
|---|---|---|
| **Cloning** | | |
| pcDNA-SARS2-N | Fwd | TACTTCCAATCCATGAGCGATAACGGCCCCC |
| | Rev | TATCCACCTTTACTGGACGCCTGAGTAGAATCGGCTG |
| pcDNA-229E-N | Fwd | TACTTCCAATCCATGGCGACCGTAAAGTGG |
| | Rev | TATCCACCTTTACTGGAATTTACTTCGTCAATGATATCAGTC |
| **CRISPR (pLentiCRISPRv2)** | | |
| IRF1 KO | Fwd | CACCGTGGTGAGAGGTGGAAGCATC |
| | Rev | AAACGATGCTTCCACCTCTCACCAC |
| **qPCR** | | |
| 229E N | Fwd | AGGCGCAAGAATTCAGAACCAGAG |
| | Rev | AGCAGGACTCTGATTACGAGAAAG |
| MX1 | Fwd | ATCCTGGGATTTTGGGGCTT |
| | Rev | CCGCTTGTCGCTGGTGTCG |
| OAS2 | Fwd | GAGTGGCCATAGGTGGCTC |
| | Rev | CGAGGATGTCACGTTGGCTT |
| TLR3 | Fwd | GCGCTAAAAAGTGAAGAACTGGAT |
| | Rev | GCTGGACATTGTTCAGAAAGAGG |
| ISG15 | Fwd | TCCTGGTGAGGAATAACAAGGG |
| | Rev | GTCAGCCAGAACAGGTCGTC |

**Supplementary Figures**

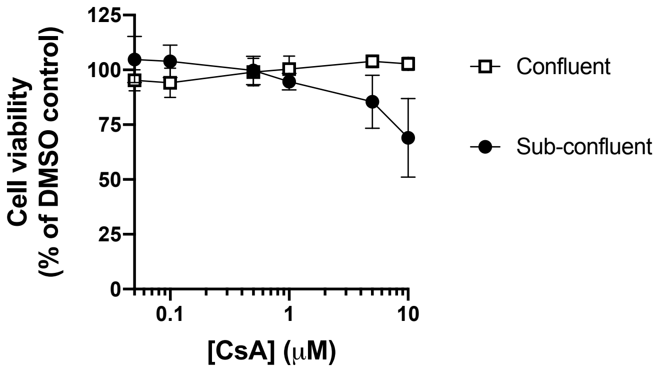


**Supplementary Figure S1. Cell viability following CsA treatment of cells seeded at different densities.** A549 cells were plated at 1 x $10^4$ cells/well (sub-confluent) or 2 x $10^4$ cells/well (confluent) in 96-well plates. The following day, cells were treated with the indicated concentrations of CsA for 48 h. Cell viability was assessed using the alamarBlue reagent and is plotted relative to DMSO control. Graph shows the average +/- SD from 2 independent experiments performed in triplicate.

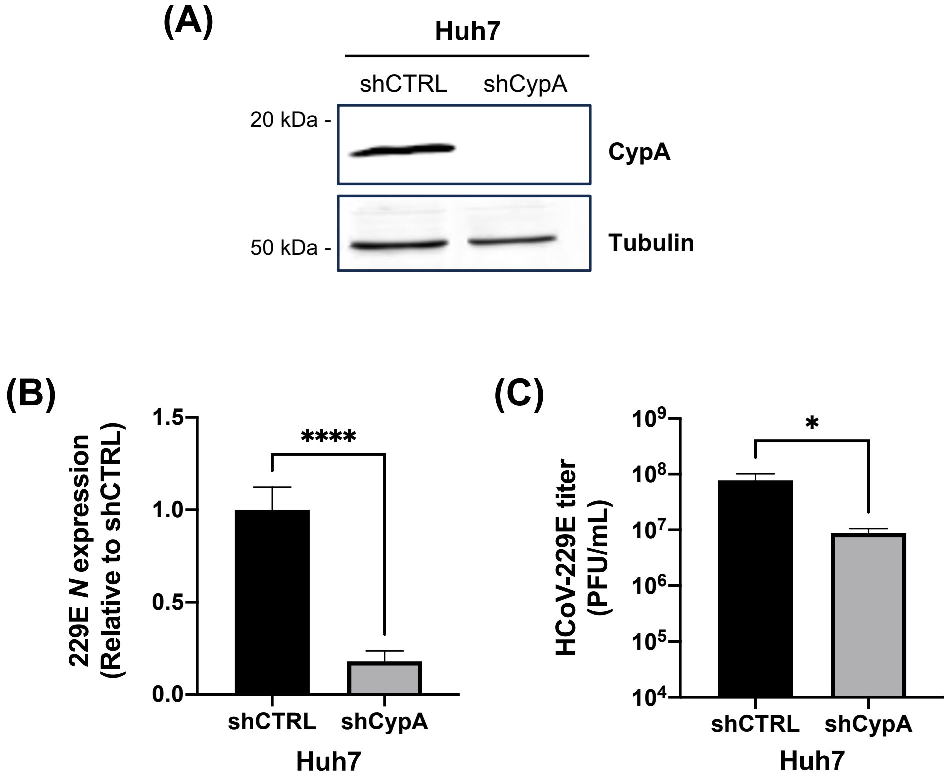


**Supplementary Figure S2. Depletion of CypA inhibits HCoV-229E replication Huh7 cells.** **(A)** Western blot confirming CypA depletion in shRNA-transduced Huh7 cells. **(B-C)** Huh7 shCTRL or shCypA cells were infected with HCoV-229E (MOI 0.01) for 2 h, at which point the inoculum was removed and the infected cells were incubated for 48 h. **(B)** HCoV-229E N gene expression was measured by RT-qPCR (normalized to actin) was reduced in shCypA cells relative to shCTRL cells. **(C)** HCoV-229E viral titer following infection of shCypA was reduced relative to shCTRL cells. Graphs show the average +/- SD from 2 experiments, with qPCR performed in technical triplicates. Significance was determined by unpaired t-test with Welch's correction (****$p<0.0001$; *$p<0.05$).

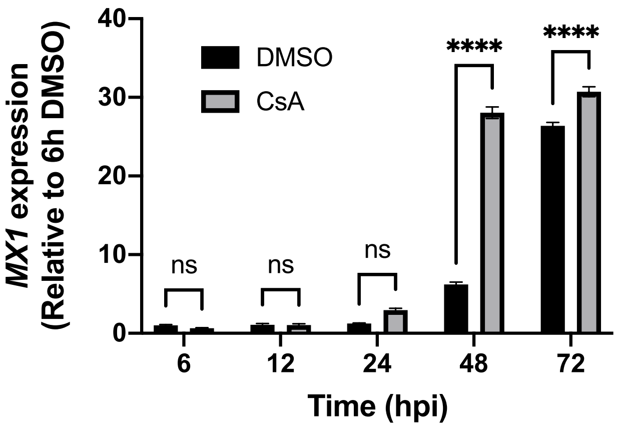


**Supplementary Figure S3. Induction of MX1 expression by CsA treatment in cells infected with HCoV-229E at higher MOI.** A549 cells were infected with HCoV-229E (MOI 0.5) and cell lysates were collected at the indicated time points to evaluate MX1 expression by RT-qPCR (normalized to actin). Graphs show the average +/- SEM from 3 independent experiments, with qPCR performed in technical triplicates. Significance was determined by unpaired t-test (****:p<0.0001, ***:p<0.001, ns, not significant).

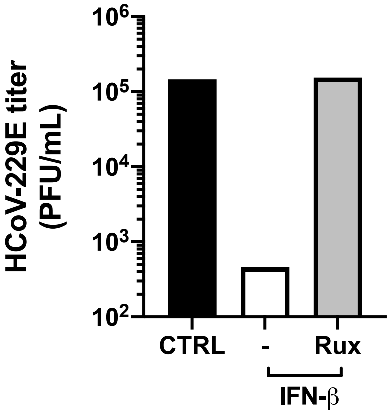


**Supplementary Figure S4. Ruxolitinib blocks the inhibitory effect of IFN-β against HCoV-229E infection.** A549 cells were infected with HCoV-229E (MOI 0.01) and were treated with recombinant IFN-β (1000 U/mL) in the presence or absence of 5 μM ruxolitinib (Rux). Viral titer was determined by plaque assay. One representative experiment is shown.

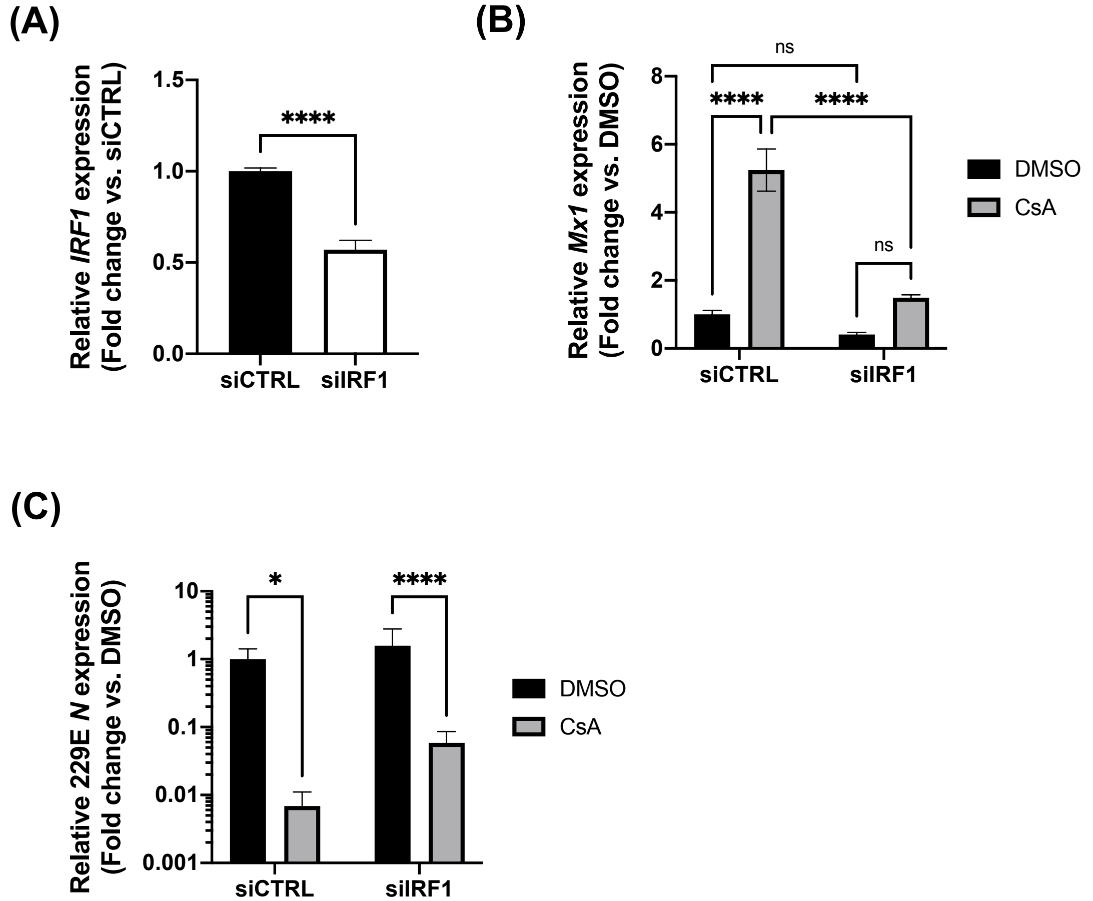


**Supplementary Figure S5. Silencing of IRF1 expression dampens MX1 induction by CsA. (A-C)** A549 cells were reverse transfected with siRNA targeting IRF1 or an siRNA control for 24 hours. **(A)** Silencing of IRF1 expression was confirmed by RT-qPCR (normalized to actin). **(B-C)** Following reverse transfection, cells were infected with HCoV-229E at MOI 0.01 and treated with 10 μM CsA or DMSO vehicle control for 48 hours. Viral N expression **(B)** and Mx1 expression **(C)** was similarly assessed by RT-qPCR. Graphs show the average +/- SEM from 3 independent experiments, with qPCR performed in technical triplicates. Significance was determined by unpaired t-test (****:$p<0.0001$, **:$p<0.01$, *:$p<0.05$, ns, not significant).
